# Optogenetic Maxwell Demon to Exploit Intrinsic Noise and Control Cell Differentiation Despite Time Delays and Extrinsic Variability

**DOI:** 10.1101/2022.07.05.498841

**Authors:** Michael P May, Brian Munsky

**Affiliations:** School of Biomedical Engineering, Colorado State University, Fort Collins, Colorado; Department of Chemical and Biological Engineering, Colorado State University, Fort Collins, Colorado

**Keywords:** Stochastic control, optogenetic, synthetic biology

## Abstract

The field of synthetic biology focuses on creating modular components which can be used to generate complex and controllable synthetic biological systems. Unfortunately, the intrinsic noise of gene regulation can be large enough to break these systems. Noise is largely treated as a nuisance and much past effort has been spent to create robust components that are less influenced by noise. However, extensive analysis of noise combined with ‘smart’ microscopy tools and optognenetic actuators can create control opportunities that would be difficult or impossible to achieve in the deterministic setting. In previous work, we proposed an Optogenetic Maxwell’s Demons (OMD) control problem and found that deep understanding and manipulation of noise could create controllers that break symmetry between cells, even when those cells share the same optogenetic input and identical gene regulation circuitry. In this paper, we extend those results to analyze (in silico) the robustness of the OMD control under changes in system volume, with time observation/actuation delays, and subject to parametric model uncertainties.

## I. Introduction

Synthetic biology seeks to create modular [2] and orthogonal [3] components to sense and actuate [4] complex logical systems [5], and which are capable of performing a variety of advanced biological behaviors [6]. Additionally, new optogenetic tools have increased our ability to actuate embedded systems within cells reliably and with strong control performance [4], [7]–[9].

Advances in so-called ‘programmed cyborg control’ have allowed for computer programmable control of cellular protein production through the use of external optogenetic inputs and smart microscopy techniques [10]–[12]. This new paradigm of digital-synthetic actuators allows for computer-modulated control of cellular systems that were previously impossible [13]. These highly adjustable systems are capable of fine tuned control without the typical drawbacks and slow timescales of chemical inputs [7]. Classical and modern control methods like PID control and model predictive control have been implemented in such systems [14], but these approaches are based on deterministic formulations that cannot allow for symmetry breaking and are therefore limited to single-input-single-output (SISO) or multiple-input-multiple-output (MIMO) control frameworks.

Because most previous control effort focus on deterministic ODE formulations of the control problem, much effort has been spent to minimize the effects of intrinsic variations (or ‘noise’) in these systems. Despite substantial work to understand and mitigate noise effects in synthetic biological systems (e.g., see the review in [15]), only a few studies seek instead to exploit noise to *improve* the control of synthetic biology processes [1], [16].

Our recent work shows that a deep understanding of noise can provide rich new control opportunities. Specifically, in [1], we find that noise, auto-regulation, and optogenetic feedback control could synergize to create a system (Fig. 1A) that systematically directs two different observed cells to two arbitrarily chosen points simultaneously, but using only a single input controller (e.g., a single-input-multiple-output or SIMO controller). Specifically, we showed that the symmetry of the OMD control problem could be effectively broken using a Fully Aware Feedback Controller (FAC) that observed all cells, a Partially Aware Feedback Controller (PAC) that observed only the primary cell, and a probabilistic Model Predictive Feedback Controller (pMPC) that observed only the primary cell and integrated a probabilistic model to represent the non-stationary probability distribution of the secondary cells in the same environment. The specification of all controllers was based on solving the chemical master equation (i.e., the forward Komogorov equation) to estimate the probability of all possible cellular states. For a simple population of one primary cell and one secondary cell, the FAC performed the best (Fig. 1C(top)), but for larger populations the FAC was computationally infeasible, and the PAC (Fig. 1C(bottom)) and pMPC controllers yielded superior results. In all cases, the controller’s knowledge and explicit treatment of intrinsic noise was critical to improve the control performance, break symmetry, and enable the SIMO control of all cells.

**Fig. 1.**
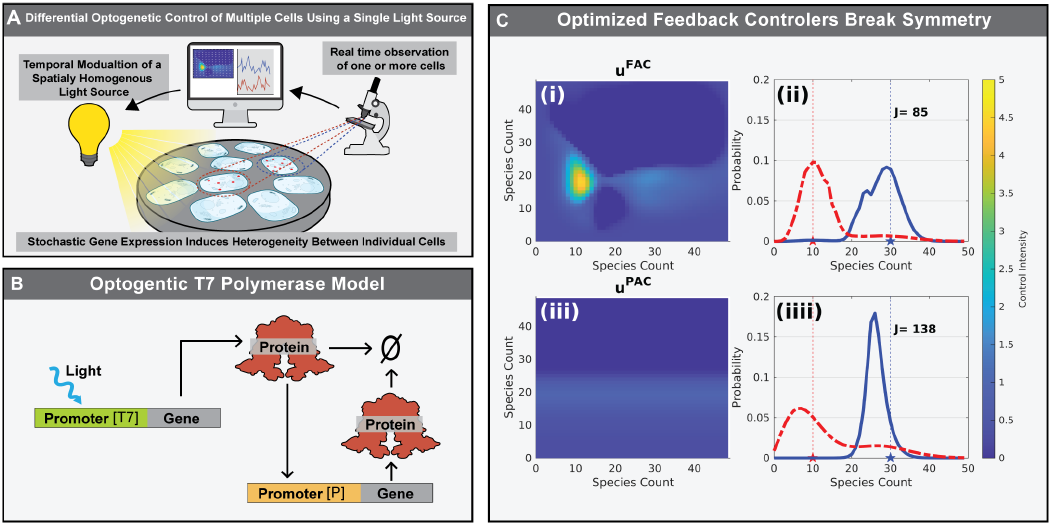
(A) Schematic of the T7 optogenetic control system (adapted from [1]). A microscope observer is partially (PAC) or fully (FAC) aware of cells; the resulting controller modulates a single spatially homogeneous light signal that perturbs all cells equally seeks to drive each genetically identical cells to a different, specified states. (B) Schematic of the optogenetically controllable T7 polymerase model, which is identical in all cells: Light activates the T7 polymerase-based production of protein, and the protein self-actuates its own production. (C) The FAC (top) and PAC (bottom) control laws for two cells (**u**^*XAC*^ (*x*_1_, *x*_2_) shown on left) break the symmetry between Cell 1 and Cell 2 and result in distinct marginal distributions for the cells’ responses (right).

In this paper, we extend the OMD control problem above to examine how the FAC and PAC controllers are affected by the cellular volume; how the control is affected by time delays in the observer or actuators; and how robust the control performance is to variations in the system parameters in individual cells. In the ‘Methods’ section, we briefly set up a stochastic chemical master equation model for multiple individual cells in the same spatially homogeneous environment, and we formulate the effects of FAC and PAC controllers on the dynamics of cells within that environment. Next, in the ‘Results’ section, we examine the effects of volume changes, time delays, and parametric uncertainties on the FAC and PAC control performance. Finally, in ‘Conclusions’ we summarize our findings and discuss potential implications that these controllers may have on future systems and synthetic biology investigations.

## II. Methods

### A. Model

We seek to analyze the dynamics of a single protein with auto-regulation and under the control of an optgentically activatable T7 polymerase (Fig. 1B). A complete model of the system is provided in [1] and consists of dimerization of production and degradation of T7 monomer components; reversible dimerization of these T7 components to form an activated T7 in the presence of UV light; reversible association of the T7 with a gene to form an active complex; T7-mediated production of the protein; auto-regulated production of the protein from a secondary self-activated promoter described by a Hill function; and degradation of the protein. This model was then simplified to a single ODE describing the protein accumulation as:

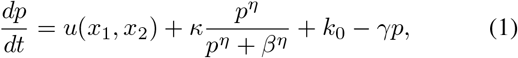

where *p* is the protein count; *κ* is the maximum auto-regulation promoter strength; *β* is the concentration at which auto-regulation reaches half strength; *η* is the auto-regulation promoter cooperativity; *γ* is the degradation rate; *k*_0_ is the promoter leakage rate; and *u* is the T7 promoter strength. The activity of the T7 promoter can be modulated in response to to the observed state, such that *u* = *u*(*x*_1_, *x*_2_) introduces the possibility of state-dependent optogenetic feedback to control the system. We note that *u*(*x*_1_, *x*_2_) refers to the effect of light on the optogenetic system rate equation, which can be converted to light intensity using the calibration curve shown in Fig. 2B in [1]. A schematic of the model is shown in Fig. 1B, and parameters were chosen to reproduce dynamics measured in [7] and are shown in are defined in Table 1 [1]. Two optimized controls laws *u* = *u*^(FAC)^ and *u* = *u*^(PAC)^ are depicted in Fig. 1C (top left and top right, respectively).

**TABLE I.** Model Parameters

| Parameter | Value | units |
| --- | --- | --- |
| $\kappa$ | 0.406 | Molecules Per Minute |
| $\beta$ | 20.0 | Molecules |
| $\eta$ | 8.00 | unit less |
| $k_0$ | 0.0001 | Molecules per Minute |
| $\gamma$ | 0.0203 | per Minute |
| $u$ | dynamic | Molecules per Minute |

**Fig. 2.**
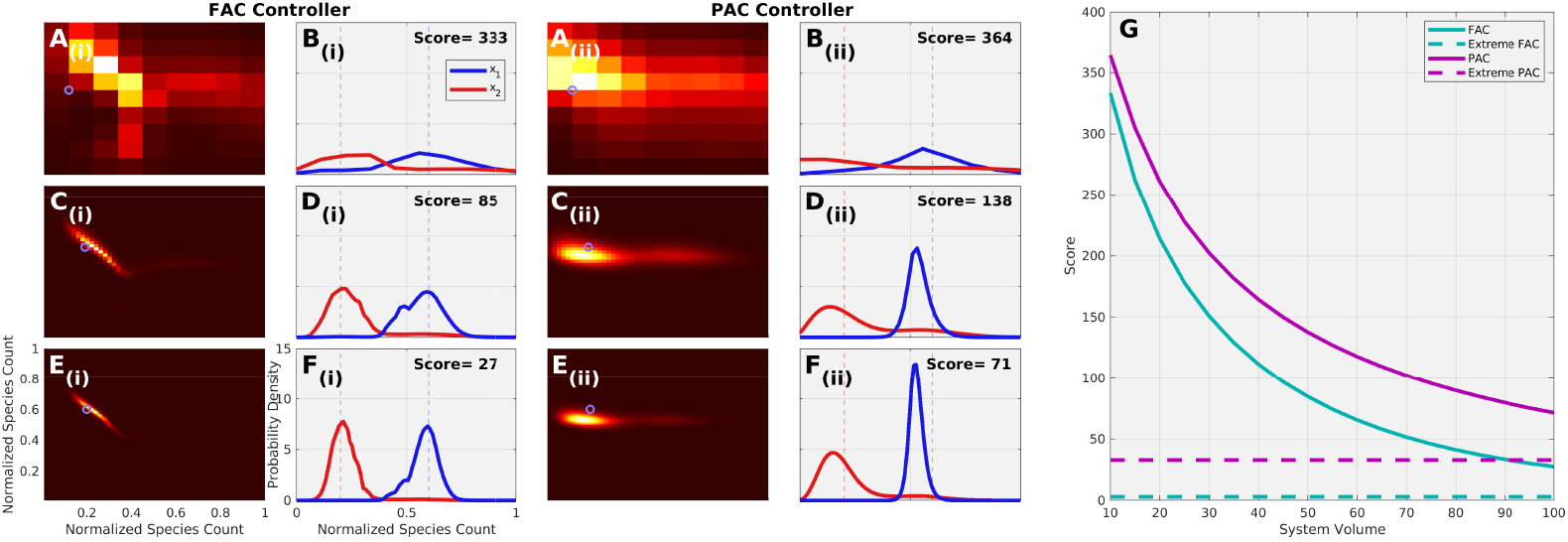
Effects of Volume Changes. (Ai-Fi) Joint (Ai,Ci,Di) and marginal (Bi,Di,Fi) probability distributions for the FAC control method at three different volumes of *N* =10 (top), 50 (middle) and 100 (bottom) relative to the target point (blue circles). (Aii-Fii) Same as (Ai-Fi), but for PAC controller. (G) FAC (cyan) and PAC (magenta) controller performance as function of system volume. Dashed lines correspond to SSA analyses at *N* =5000. In all analyses, the system model and PAC and FAC controllers are unchanged from [1] other than their adaptation to the new system volume size.

### B. SSA and Finite State Projection Analysis Methods

To recast the above ODE formulation into the discrete stochastic representation, we define the enumerated *i* = 1, 2, … states of the system as the tuple of the numbers of proteins in each cell as 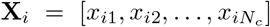, and we define the stoichiometry vectors as the change in state following one reaction event (e.g., **X**_*i*_ *→***X**_*i*_ + **s**_*ν*_), where the 2*N*_*c*_ possible reactions are:

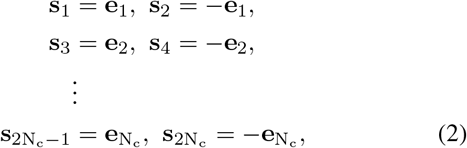

where each 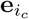 corresponds to the Euclidean basis vector for the 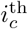 cell. The corresponding propensity functions are:

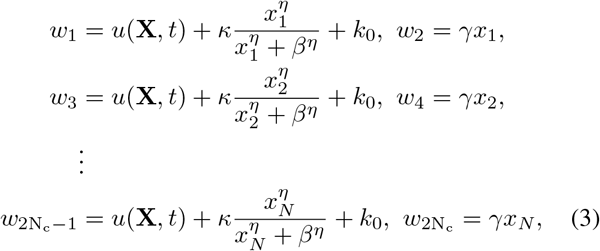

With these stoichiometry and propensities, we can formulate and run the Gillespie stochastic simulation algorithm [17], [18] to generate representative trajectories of the stochastic process.

For the same specifications of the stoichiometry and propensity functions, the chemical master equation can also be formulated in matrix/vector form as:

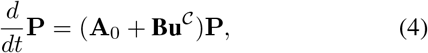

where **P** = [*P* (**X**_1_), *P* (**X**_2_), …]^*T*^ is the enumerated probability mass vector for all possible states of the system; **A**_0_ is the infinitesimal generator of the stochastic process due to the autoregulation promoter and degradation events; **u**^𝒞^ = [*u*^𝒞^(**X**_1_), *u*^𝒞^(**X**_2_), …]^*T*^ is the collection of state-dependent control inputs associated with each state; and **Bu**^𝒞^ is the infinitesimal generator for the T7 promoter expression by which the effects of controller 𝒞 are introduced into the process.

The generator **A**_0_ can be constructed according to:

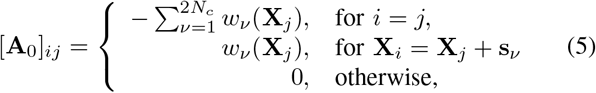

and the infinitesimal generator **Bu**^𝒞^ of the controller is given by

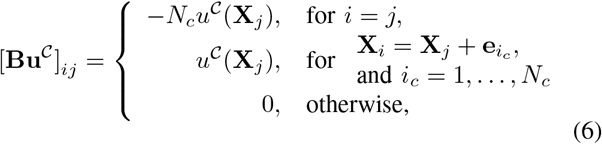

where *u*^𝒞^(**X**_*j*_) is the specification of the 𝒞 controller in terms of the (partially observed) state.

For a given controller, the steady state of the system (**P**^*∗*^) found by solving Eq. 4 is given by

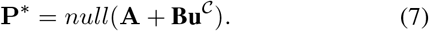

### C. Controller Performance Ranking and Optimization

A performance metric is specified in order to compare the outcomes of different controllers and to define an optimization for the control law. Specifically, we define the performance score (*J*), as the expected steady state Euclidean distance of the process from a specified target state according to *E* {(**X** *−***T**)^2^}. This score can be rewritten in terms of a precomputed set of linear weights (**C**) and the steady state probability distribution (**P**^*∗*^ from Eq. (7)) given by

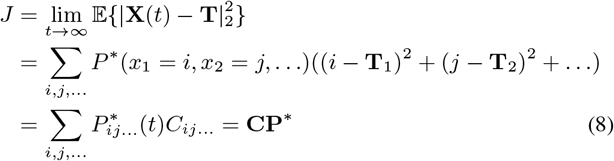

Two controller designs, a fully aware controller **u**^*FAC*^ = **u**^*FAC*^(*x*_1_, *x*_2_) and a partially aware controller **u**^*PAC*^ = **u**^*FAC*^(*x*_1_), were specified in previous work [1]. The FAC observes both *x*_1_ and *x*_2_ when making its control decision, while the PAC only uses information from *x*_1_ to derive its control input. Each was optimized to drive the OMD control problem with high probability to a target state of **T** = [30, 10]^*T*^. We note that despite the fact that the PAC only uses information from *x*_1_, the control input is capable to control both *x*_1_ and *x*_2_ almost as well as the FAC controller (Fig. 1C, compare top and bottom right). As defined in Eq. 6, the control matrix **Bu**]^𝒞^ changes depending on the control type used (**u**^*FAC*^ or **u**^*PAC*^). For the purposes of this work, we adopt the precomputed values for the **u**^*FAC*^ and **u**^*PAC*^ controllers (shown in Fig. 1C, top and bottom left), and we leave these unchanged as we perturb the system to test how the performance of the controller depends on the volume of the system, time delays in the observation or actuation of the control signal, and perturbation to the system parameters.

### D. Rescaling for Different System Volumes

To examine how the system size (or volume) affects the ability of the controller to force the state to the target value, we introduce a volume scaling parameter *α* = (new volume)/(original volume) that scales the state (**X** = [*x*_1_, *x*_2_]) whose dimensions depend on molecule numbers (or concentrations). As a result of this scaling, parameters remain the same, but the rate equation as a function of state becomes *w*_*µ*_(**X***/α*). In order to use the same control law or performance metric as for the original system size, the state must also be scaled by 1*/α* before computing the appropriate functions of that state, e.g., *u*^𝒞^ = *u*^𝒞^(**X***/α*) and

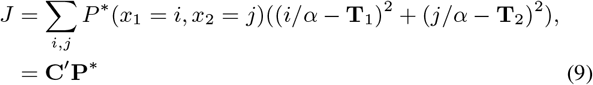

Finally, as the original controllers **u**^*FAC*^ and **u**^*PAC*^ were only defined for integer numbers of molecules, fractional values are determined using bi-linear interpolation.

After rescaling, the SSA analysis is conducted using Eq. 2 and the modified version of Eqs. 3, and the truncated CME analysis of the model is found by substituting the scalable rate equation into Eqs. (5) and (6). We note that the original system volume was proportional to *N* = 50, such that the truncation size of the CME in Eq. 4 for a two cell system led to infinitesimal generators **A**_0_ and **Bu**^𝒞^ of the size 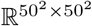.

### E. Introduction of Observation/Actuation Time Delays

To test the effects of time delay, a time-delay SSA was developed by recording the piecewise constant recent history of the system and calculating the current light at time *t* in the following delayed manner

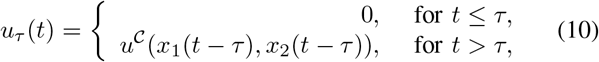

where *τ* is the time delay of the observation or actuation, and *u*^𝒞^ is the previously optimized control law for **u**^*FAC*^ or **u**^*PAC*^. We note that the time delay stochastic process was only simulated using the SSA because an appropriate direct FSP/CME integration procedure is not known.

### F. Extrinsic Parameter Perturbations

To estimate the effects of inaccurate models or variations of parameters in different cells, a parameter perturbation analysis was performed by taking each individual parameter *λ* ∈ {*κ, β, k*_0_, *η, γ*} and multiplying that parameter by a positive scalar value ranging from 1*/*2 to 2. We examine two cases for such perturbations: First, to analyze the effect of different parameter perturbations in different cells (e.g., extrinsic variability), parameters of one cell would be held constant while the parameters of the other cell is modified. Second, to examine the effects of a global error in the model that affected both cells, we would perturb the same parameter for both cells simultaneously.

## III. Results

### A. Effects of Changes to Volume Size

It is well known that systems are intrinsically more noisy when the numbers of important molecules are small, and systems are less noisy when the number of molecules are large. Moreover, we know from [1] that the existence of intrinsic noise is essential for the realization of the OMD control strategy and its control performance. To better understand the effect of intrinsic noise due to molecule count on control performance, we re-scaled the system to have to same dynamics over a larger (or smaller) system sizes, and we assess the control performance at these different volumes.

To explore how the size of the system volume changes the intrinsic noise level and the control performance, we solved the FSP steady state for the volume-adjustable system described above. Specifically, we changed the volume (originally *N* =50 in [1]) from *N* =10 to *N* =100 in steps of five, and at each point the score of the system is calculated using the FSP at steady state. Figs. 2 (Ai - Fi) show the joint (left plots) and marginal (right plots) distributions of the two cells relative to the specified target position (circles in joint distributions and vertical dashed lines in marginal distributions) for the FAC controller and for volumes of N = 10 (A,B), 50 (C,D) and 100 (E,F) cells. As the volume changes from N=10 (Fig. 2, Ai and Bi), to N=50 (Fig. 2, Ci and Di), to N=100 (Fig 2, Ei and Fi), the distributions become much better centered around the intended target values, and the performance score improves from *J* =333 to 85 to 27, respectively. The PAC controller performance also improves considerably with volume as shown in Figs. 2 (Aii - Fii).

Fig. 2G shows the trend of the performance score versus the volume size for both the FAC (solid cyan line) and the PAC (solid magenta line) controller. This improvement in performance appears to approach a small value as the volume size goes to infinity, but since the size of the FSP increases with the square of the system volume, systems much larger than *N* =100 (where 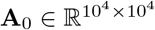) are difficult to calculate. To bypass this limit in the FSP, we used a series of Gillespie SSA to simulate a system with a much larger volume of *N* =5,000 molecules. The performance score estimates of this large volume SSA using the FAC and PAC were 2.9 and 33 respectively, which are plotted as dashed lines in Fig2G. We note however, that systems with larger volumes take longer to reach a steady state distribution, and it is unclear if performance improvements can be obtained with further increases to the system volume. However, for all volumes considered, we found that both controllers monotonically improved with increased volume and that **u**^*FAC*^ always outperforms the **u**^*PAC*^ at any system volume.

These data have a couple important implications for the design of controllable synthetic systems as well as for the determination of a usable control algorithm. First, they suggest that increasing system volume increases steady state control performance even if the controller itself depends on noise for its successful implementation. Second, we showed that designing a controller to work for a system of one volume size can result in a controller that works even for another much larger volume. This second result has crucial practical significance, because it is relatively easy to search over a control law defined on an *N × N* grid of states when the size is relatively small (e.g., when *N* =50 as in [1]), but to design the controller for a larger system (e.g., for when *N* =5000 as simulated with SSA above) may be computationally prohibitive. This result implies that it may be possible to optimize controller using coarse-grained FSP analyses for small volume problems and adapt these via interpolation for use with larger, more realistic systems, that exceed the computational limit of standard FSP computations.

### B. Effects of Time Delays on Controller Performance

Time delay is a common occurrence in practical control applications that usually degrades performance and may even produce oscillations or destabilize a physical system. We hypothesized that time delays in the observation or control signal actuation would adversely affect the ability to control the individual cells due to their respective target states. To test the effect of time delay, we analyzed the performance of the original FAC and PAC controllers using the time delay Gillespie SSA, each considering time delays ranging from from 0.01*/γ* to 10*/γ* (where the residence time 1*/γ* sets the dominant time scale of the protein concentration).

Fig. 3 shows the effects of time delay using the two controllers, with panels Ai - Fi showing results for the FAC controller and panels Aii-Fii showing results for the PAC controller, and panel G showing the score of the controllers versus the time delay. From the figures, it is clear that performance is rapidly degraded as the delay approaches and then exceeds the characteristic time scale of the process. At low time delays (below *τ* = 0.07*/γ*), the FAC outperforms the PAC (*J* =87 versus 146 at *τ* = 0.01*/γ*) but at moderate time delays (above *τ* = 0.07*/γ*) the PAC outperforms the FAC (*J* =170 versus 219 at *τ* = 0.1*/γ*)). This data taken together show that time delays much larger that 0.07*/γ* is detrimental to the FAC controller; delays beyond 0.2*/γ* are detrimental for the PAC controller; and the best choice in controller is dependent the level of time delay in the system. For extreme levels of time delay, both systems lose their asymmetry, and their scores become much worse (*J* =1041 and 931).

**Fig. 3.**
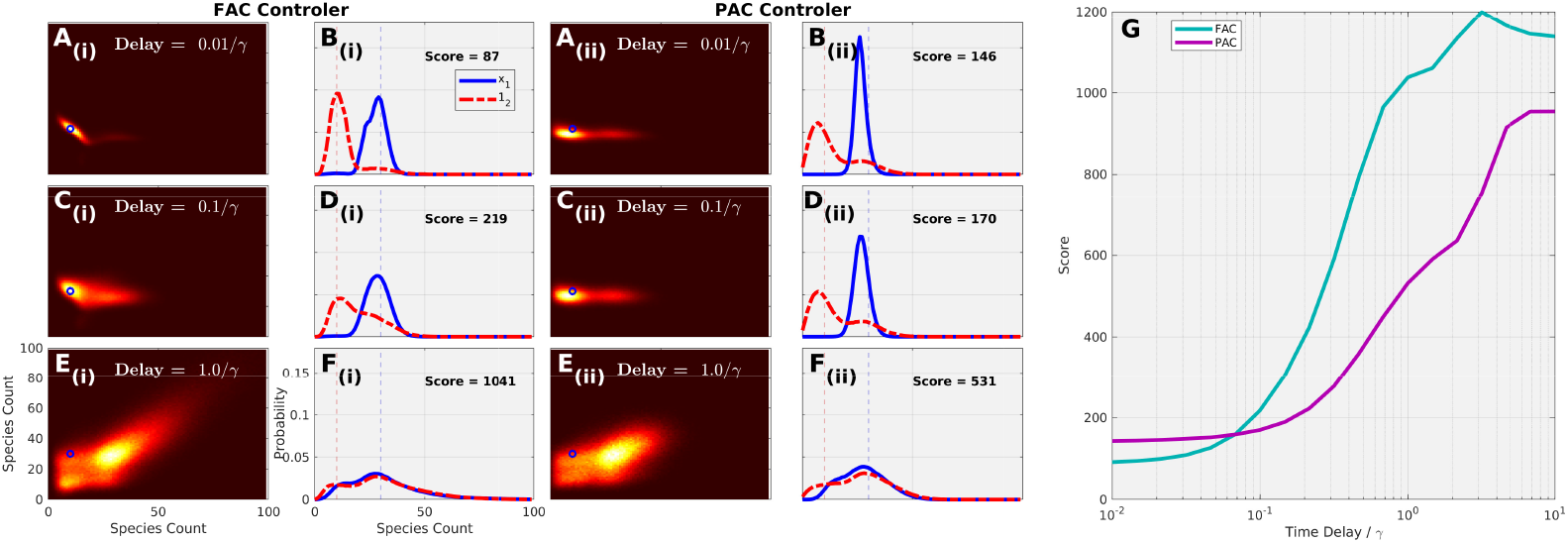
Effects of time delays. (Ai-Fi) Joint (Ai,Ci,Di) and marginal (Bi,Di,Fi) probability distributions for the FAC control method at three different time delays of *τ/γ*=0.01 (top), 0.1 (middle) and 1 (bottom) relative to the target point (blue circles). (Aii-Fii) Same as (Ai-Fi), but for PAC controller. (G) FAC PAC performance as function of time delay show that PAC outperforms the FAC when time delay is greater than 0.07*/γ*. In all analyses, the system model and PAC and FAC controllers are unchanged from [1].

### C. Effects of Parameter Errors or Extrinsic Uncertainties

It is rare that any parameters are perfectly known (especially for biological processes), and real systems often exhibit slightly different parameters due to extrinsic variations. We hypothesized that perturbations of cell parameters would influence the control performance of the OMD controller. To test this, we perturbed individual parameters of Cell 1 while holding the parameters of Cell 2 constant. The data for such perturbations are shown in Fig 4(Ai-Ei). The analysis was then repeated, this time holding the parameters of Cell 1 constant and perturbing the parameters of Cell 2 (shown in Fig. 4(Aii-Eii)). Finally, both Cell 1 and Cell 2 had their parameters jointly perturbed by the same amount (Fig 4(Aiii-Eiii)).

**Fig. 4.**
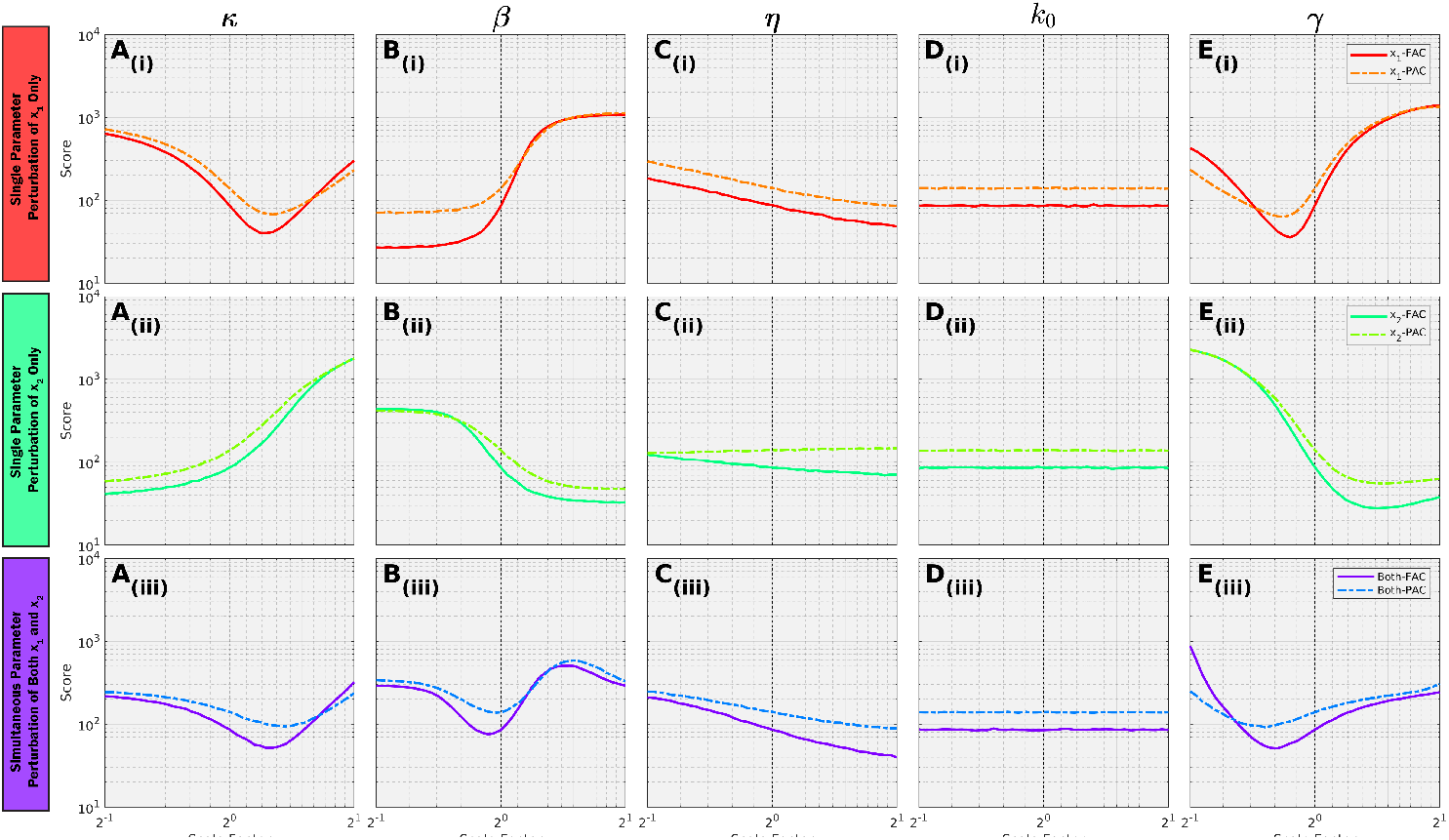
Effects of parametric variations to Cell 1 (top row), Cell 2 (middle row) or to both cells simultaneously (bottom row). Each column show the effect of changing a single model parameter (e.g., *κ* in the leftmost row). Trends for the *FAC* (solid lines) and PAC (dashed lines), but the optimal controller can change under parameter variations. In all analyses, the system model and PAC and FAC controllers are unchanged from [1].

Considering perturbations to one cell at a time (Fig. 4 top two rows), we found that perturbations to different parameters had different effects. For example, Fig4(Bi) shows that increasing *β* in Cell 1 worsens performance while increasing *β* in Cell 2 improves performance, showing that fortuitous asymmetry in the cells may lead to better control performance. In some cases, these effects were not monotonic; for example increasing *κ* in Cell 1 is highly advantageous up to a limit after which the control performance degrades rapidly. In other cases (such as for *k*_0_), the effect of perturbations on performance is insignificant even for relatively large (50%) perturbations. When parameters of both Cell 1 and Cell 2 are jointly changed (Fig. 4 bottom row), some parameters (e.g., *κ, γ*) reached minima that performed better than the original, which suggests that the physical system itself has room for optimization beyond the original design that could lead to better control performance, *even without changing the control law*.

When examining the effects of changing parameters when using the **u**^*FAC*^ (solid lines) or the **u**^*PAC*^ (dashed lines), general trends typically remained the same in that changes to a parameter which caused a decrease in the control performance of the FAC also tended to decrease the performance of the PAC. However, these effects were not always equal (e.g., for large *κ* values or small *γ* values) and the simpler PAC occasionally outperforms the FAC controller.

## IV. Conclusion

The modeling of noise is known to be critical to the success of synthetic biological systems. Our findings show that noise can provide beneficial aspects to process controllability by breaking symmetry and enabling a single controller to simultaneously module the behavior of multiple different cells to reach distinct phenotypes. We show that such an Optogenetic Maxwell’s Demon control strategy can drive cells to different stable points despite moderate external influences, such as volume changes to the system, time delays in the controller and variations in the system parameters. These analyses also show that approximate models can be used to design effective controllers, and that systems and controllers can be jointly optimized to improve performance. Given the relative robustness to time delays exhibited by the PAC controller, we believe that more sophisticated probabilistic model predictive controllers, such as that introduced in [1], could be implemented in future work to predict effects of time delays and create controllers that are even more robust to time delays. Additionally, we believe that the performance score in the control design could be extended to account for a parameter uncertainties and a parameter-robust controller could be developed to sacrifice some performance at baseline parameter combinations but be better able handle a range of parameter combinations.

## Notes

Research reported in this publication was supported by the National Institute of General Medical Sciences of the National Institutes of Health under award number R35GM124747.

### Competing Interest Statement

The authors have declared no competing interest.

